# A systems genetics approach to uncover mitochondrial drivers of heart failure reveals mitochondria-nuclear cross talk in genetically diverse mouse strains

**DOI:** 10.64898/2025.12.12.693865

**Authors:** Sriram Ravindran, Todd H. Kimball, Brian Gural, Caitlin Lahue, Christoph D Rau

**Author notes:** Corresponding author: (CDR).

## Abstract

The contribution of mitochondrial genetics to heart failure is thought to be reciprocal. Its complex traits are influenced by genetic and environmental factors. Genetically diverse mouse strains form a vital repository to uncover the interaction between the mitochondrial and nuclear genomes underlying heart failure due to their traceable genetic origins, maternal lineages and controlled genetic variation. Using systems genetics, we studied this cross-talk in the Collaborative Cross (CC) mice challenged with heart failure (HF).

We used 63 strains of CC-mice that grouped into 8 mitochondrial haplotypes and subjected them to HF using isoproterenol (Iso), a beta-adrenergic stimulant that mimics progressive stress induced HF in humans. The Alzet osmotic pumps delivered a consistent dose of the drug for 21 days following which the mice were euthanized for organ collection. A group of mice with saline loaded pumps acted as control (Ctrl). Baseline and end of study echocardiography were recorded for these mice. Bulk RNA sequencing was carried out on the left ventricles and data analyzed on R-studio.

The CC-strains showed differences across the haplotypes for organ weights and heart function. Noticeable treatment specific gene expression differences were observed for 34 nuclear encoded genes from MitoCarta3.0 unlike mt-DNA encoded genes that were insignificant after correcting for sex and haplotypes. These genes were associated with multiple metabolic pathways in the cell. Trait-GWAS associations with markers from the mitochondrial genome were observed only for mice from Iso group with cell surface area (p = 6.19e-06) and change in ejection fraction (p = 9.22e-05) as top hits under FWER threshold of 0.01. *Cyfip2* was the top candidate gene (p = 2.66e-07) among the 831 hits (47-MitoCarta & 784 other nuclear) based on our trans-eQTL analysis. The eQTL genes influenced critical pathways of OXPHOS, myogenesis, apoptosis etc. Our approach uncovered 24 gene candidates associated with mt-DNA and HF that overlapped with our previous mi-eQTL reports. CC-mice revealed ancestry dependent effects underlying HF for studying mito-nuclear interactions. Despite establishing differences, modeling these interaction needs development of complex cybrid systems to evaluate the bidirectional impact on HF.

**Author Summary:** The Collaborative Cross (CC) mouse is a genetically diverse population that has been used to study complex diseases. Recently, our group has comprehensively characterized the heart from 63 strains of the CC and reported genetic associations with heart failure (HF). Traditionally, genetic abnormalities underlying a disease are related to the nuclear genome but there exists an alternate genome within the cells, the mitochondrial DNA (mt-DNA), whose contribution is understudied. The mt-DNA of CC mice is maternally derived from the 8 founder strains, and our sequencing data holds this information, allowing us the ability to use them to study the contribution of mitochondrial ancestry (haplotypes) to HF. We found haplotype differences in terms of cardiac function and gene expression (both nuclear and mitochondrial) that were associated with HF. It revealed stress induced changes in the heart (trait and gene expression) linked with regions in the mt-DNA that encode genes involved in oxidative phosphorylation and metabolism. We uncovered 24 high-confidence nuclear gene candidates that agreed with our previous analyses and were associated with regions on the mt-DNA. These findings open up new opportunities to study the CC and advance the understanding of Mito-nuclear interactions in HF pathophysiology.

## Introduction

Heart failure (HF) affects 55.5 million people globally, yet the complexity of the disease, with many inciting factors, a spectrum of presentations and numerous avenues of progression has made the design of therapeutic options for reversing HF progression difficult(1). Central to cardiac function is the mitochondrion, which occupies a third of the volume of the adult cardiomyocytes and, in its role as the ‘powerhouse of the cell’s supplies ∼95% of heart’s ATP. Growing evidence suggests that mitochondria have a more nuanced role than merely a ‘powerhouse’ and are, instead, crucial processors of cellular signals through the mitochondrial information processing system, or MIPS (2). Consequently, there is increasing interest in how mitochondria regulate signal transduction (PMC9960245), yet the role of genetics on mitochondrial signaling in the context of HF remains poorly understood(3) (4).

Prior work in humans has shown associations between mitochondrial haplogroups and global mt-DNA content with coronary artery disease (5, 6). Similar research in HF has proven difficult due to greater challenges in obtaining tissue samples and regulating environmental variability. These challenges lead to reduced power and limited ability to perform genetic scans such as GWAS on mt-DNA variants in HF. Our group has a history of using mouse genetic reference populations (GRPs) to circumvent these challenges(7–14). GRPs offer stable, reproducible platforms upon which to perform powerful systems genetics approaches.

The Collaborative Cross (CC) is a murine genetic reference population (GRP) in which eight highly diverse inbred lines (129S1/SvImJ, A/J, C57BL/6J, NOD/ShiLtJ, NZO/HiLtJ, CAST/EiJ, PWK/PhJ, and WSB/EiJ) were mated together in a controlled 8-way breeding funnel to produce a number of new inbred lines that blend the genetic contributions of those eight founders (15–17). The CC holds promise in exploring mitochondrial signal transduction due to their diverse maternal ancestry, enabling high mapping resolution for studying genetic associations. Better understanding how heritable mitochondrial changes affect the pathogenesis of HF would be a crucial leap forward in our understanding of HF and the role of mitochondria in its progression. This is particularly true for the CC, where identical mitochondrial haplotypes are paired with diverse nuclear genomes, offering an ideal platform for complex trait discovery.

We recently completed a study of HF progression in 63 CC lines and the 8 CC founder strains after treatment with the beta-adrenergic agonist Isoproterenol (18). We noted significant phenotypic variability both at baseline and after catecholamine overdrive and identified a number of genome-wide significant loci linked to HF-associated phenotypes. Like many GWAS studies, our analysis did not extend to an analysis of mitochondria. mt-DNA GWAS is often overlooked for a variety of reasons – a lack of mt-DNA sites on genotyping chips, the presence of heteroplasmy (variation in mt-DNA sequence *within* a cell), and simply the difficulty of isolating both the 16kb mt-DNA and the 10s-100s of mb per chromosome nuclear DNA from the same samples. We have previously provided brief insights into how to approach GWAS studies to explore mitochondrial genes in order to identify causal variants for HF in both nuclear and mitochondrial genes (19).

Consequently, in this study we apply these insights to the mice of the CC. Due to the way in which the CC was constructed, we know without genotyping the mitochondrial backgrounds of each strain as they represent one of 8 different mitochondrial haplotypes each arising from one of the 8 founder lines of the CC. We test the hypothesis of whether strain-specific mitochondrial gene expression profiles within these haplotypes promote susceptibility to HF. Our approach integrates RNAseq data to characterize mt-DNA variations with gene expression patterns (mitochondrial and nuclear) in the context of HF phenotypes. Our analysis uncovers sites on the mt-DNA that either affect or are affected by both nuclear gene expression and cardiac phenotypes. By examining these mito-nuclear interactions (crosstalk between both the genomes), we provide a framework for crucial systems-level understating of mitochondrial disease biology that is often obscured in traditional studies (20).

## Methods

### Ethics statement

All the mice experiments were conducted with approval from the Institutional Animal Care and Use Committee (IACUC) at the University of North Carolina, Chapel Hill, NC, USA. All experiments were conducted according to ARRIVE guidelines.

### Study design

We used 63 strains of CC mice and 8 founder lines and categorized them into 8 mitochondrial haplotypes (A-H) (Supplementary table 1) based on their maternal lineage through the breeding funnel (17). The mice were maintained in individually ventilated cages (Tecniplast,USA) under controlled environments at UNC. Mice aged 8-10 weeks were utilized for experiments.

Briefly, heart failure was experimentally induced in these mice through sustained delivery of Isoproterenol (30 mg/Kg body weight/day) using subcutaneously implanted Alzet Osmotic pump (21). The mice were divided into two groups control (Ctrl) and isoproterenol (Iso). The control group received saline loaded pumps. At 9 weeks, the mice underwent minor surgery at the Cardiovascular Phenotyping core (McAllister Heart Institute, UNC, Chaple Hill) to receive the pumps (day 0) and monitored for signs of discomfort until the end of study (day 21). The body weights and echocardiography data were recorded under anesthesia on a VevoF2 ultrasonic imaging system (Visual Sonics) before implanting the pumps that served as baseline and again on day 21. Images were captured in B-Mode and M-mode to measure cardiac physiology, and measurements were carried out on VevoLAB. At the end of study, mice were euthanized by isoflurane drop method followed by organ collection and weighing. Tissue samples were stored at −80 °C for further analysis. For more information see our manuscript in which we describe the CC-HF cohort in greater detail (18).

### RNA sequencing

Heart samples were thawed to extract RNA from left ventricular tissue using the QIAGEN RNA-Mini Kit (Cat: 74106). Samples having 260/230 > 1.6 and RIN > 7 were included for analysis. A total of 440 samples were used for library preparation using five 96-well Mercurius Full Length mRNA BRB-seq kits from Alithea Genomics, MD, USA (Cat: 10513) per manufacturer’s instructions. Reads were analyzed on a single Novaseq X 25 Gb chip according to the Alithea protocol.

### Data processing and analysis

#### QC and mapping

The raw reads were processed as per the pipeline instructions from Alithea genomics. Following a FastQC check, samples were separated based on their unique barcodes, followed by alignment and mapping to the mouse reference genome (GRCm39) using the STARSolo algorithm (STAR v2.7.9a). Resulting gene-count matrices generated for each plate were merged in R and sample metadata was added to create a DESeq object for further analysis.

#### Differential gene expression (DGE) analysis

The DESeq2 R-package (22) was used to model gene expression difference across haplotypes considering the sum of effects of the plate number (a potential batch effect), treatment (Ctrl/Iso), and sex. The count data was subset to a list of genes that were encoded by the mitochondria (mt-‘gene’) and those that were translocated to the mitochondria as listed in the MitoCarta3.0 database, separately for DGE on the two data sets (23). Genes were retained for analysis if they had at least 10 counts in 50% or more of the samples ensuring sufficient expression across the dataset.

#### Mitochondrial markers

The marker positions (loci) from the GigaMUGA SNP arrays for the CC founders (https://kbroman.org/qtl2/pages/move_to_build39.html) were downloaded for the mitochondrial genome and the genotypes of CC-strains were imputed based on founder haplotypes. We tested the genome wide associations for 21 marker positions as listed in the supplementary table 2).

#### Trait-GWAS

We tested the association of cardiac traits with mitochondrial loci using a linear model which models phenotypes based on founder haplotypes and estimates a haplotype-trait association. Traits were z-transformed and corrected for sex and cross-experimenter biases and associations found for both control and Iso groups independently. The genome-wide significance threshold (P=0.0029) was set based on Family-Wise Error Rate (FWER) of < 0.01 as originally described (24).

#### Trans-eQTL

We defined trans-eQTL (expression Quantitative Trait Loci) as loci within the mitochondrial genome significantly associated with nuclear gene expression. We adopted the same testing approach using a linear model as above to select the significant nuclear gene targets that influence or are influenced by loci in the mitochondrial genome.

#### GSEA for hallmark pathways

The Cluster Profiler R-package (25) was used for an assessment of Hallmark pathways influenced by the genes that were differentially expressed and trans-eQTL hits. A ranked and sorted gene list based on the test statistic and p-value was used to predict hallmark pathways with an adjusted p-value cutoff of 0.05. The custom pathways for mitochondrial genes were imported from MitoCarta3.0 for testing genes that were differentially expressed under Iso stress. In order to capture strong haplotype-specific gene expression signals that were obtained from DGE analysis, the maximum absolute score of ranks for the genes was considered for GSEA, while median ranks were used for the trans-eQTL hits as they were better related to the associated mitochondrial SNP positions.

#### Overrepresentation testing

Ranked list representation analyses were done to verify if eQTL hits were overrepresented within MitoCarta genes for Ctrl and Iso results. A Wilcox rank-sum test was executed on the data to verify if the rank distribution for the genes differed significantly (p <0.05) for Iso vs Ctrl.

#### Hierarchical clustering

The dendextend R-package (26) was used to construct dendrograms for haplotype level data. We compared hierarchical clustering of traits from the 8 haplotypes to their founder mt-DNA sequences for the two treatment groups using tanglegrams for an assessment of the entanglement scores (ES). The tanglegrams inform whether the strains with similar mt-DNA sequence have similar traits. A score of 0 indicates high similarity (meaningful relationship) between the two dendrograms while 1 indicates the opposite. Additionally, the cophenetic correlation coefficient (CPCC) was used to quantify the degree of similarity. The CPCC ranges from −1 to 1, with values closer to +1 indicating preserved pairwise distance or a reliably comparisons of two dendrograms, and values close to −1 indicate an inverse relationship. A CPCC of 0 indicates no relationship between the dendrogram structure and the data distances. Similar comparison was made for mitochondrial gene expressions from the CC-strains to their founder strains individually for the two treatment groups and among the CC-strains (Iso to Ctrl) and their founders (Iso to Ctrl) to understand the degree of similarity of gene expressions across the haplotypes.

#### Data availability

Phenotypic data from the CC-Heart Failure Project can be found on Mendeley Data: https://data.mendeley.com/datasets/vyf5x4ygrv/1. The transcriptomic data is deposited on the Sequence Read Archive #*PRJNA1331689*.

## Results

### Grouping mice by mitochondrial haplotype reveals trait differences linked to cardiac performance

The strains of the CC are derived from 8 founder lines, and their breeding funnels are recorded, meaning that we can uniquely link the mitochondria of CC strain to one of the original 8 mitochondrial haplotypes (See Supplementary table 1). Each haplotype group was split based on treatment (Iso vs Ctrl) and phenotypic differences across static (organ weights, %fibrosis and cell surface area) and dynamic (Echocardiography parameters) traits were analyzed for each haplotype (Fig. 1a and 1b).

**Fig. 1.**
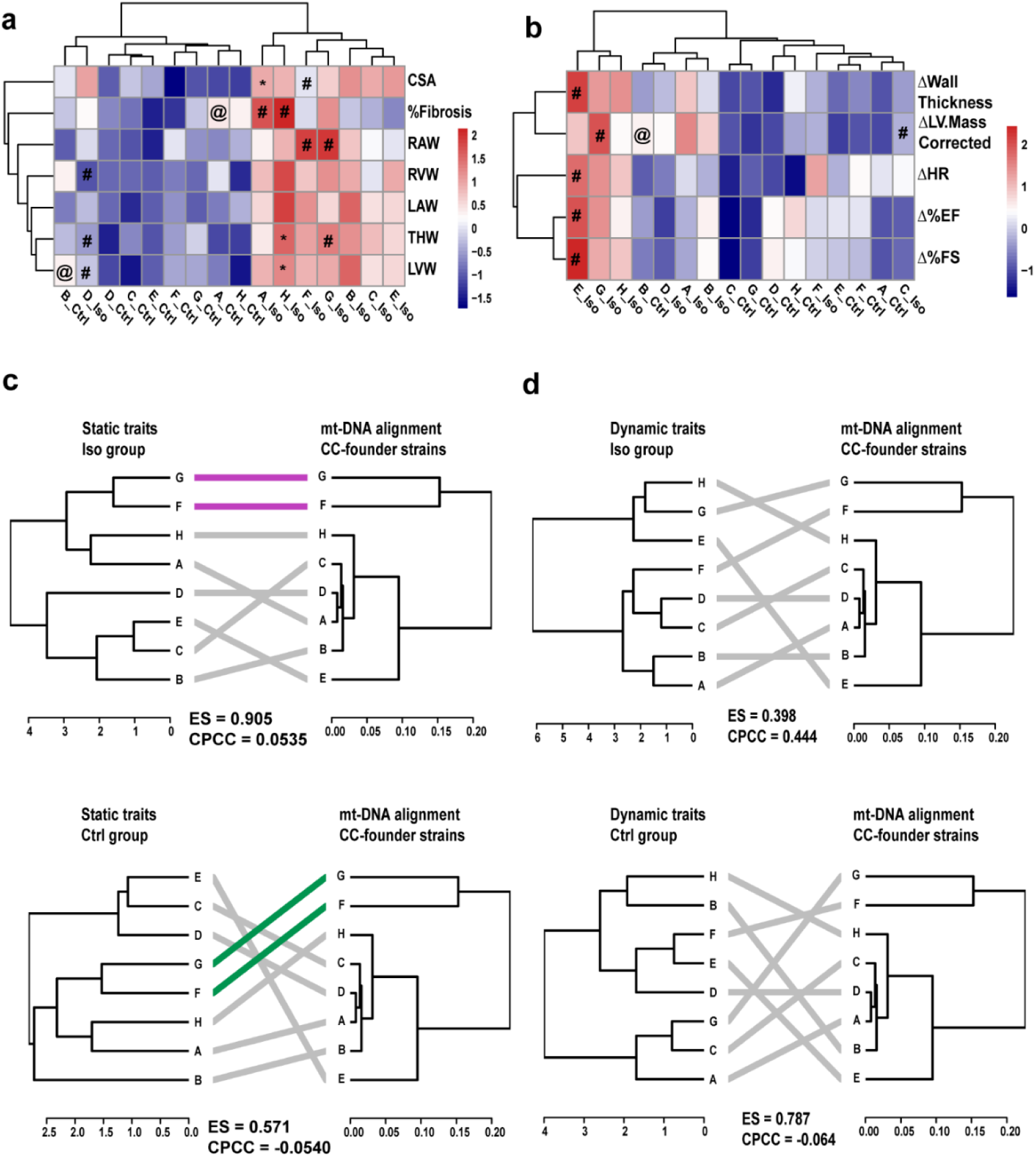
Observed trait differences across the 8 mitochondrial haplotypes for the two treatment groups (Ctrl and Iso) are shown as heatmaps for a) static traits and b) dynamic traits. Tanglegrams representing association between the hierarchical clustering of traits from the CC-haplotypes and mt-DNA sequences from the CC-founder strains are displayed for c) static traits; top – Iso, bottom-Ctrl and d) dynamic traits; top-Iso, bottom-Ctrl. Significant (p < 0.05) for * Iso vs Ctrl # vs all other haplotypes in Iso, @ vs all other haplotypes in Ctrl. ES- entanglement score, CPCC-cophenetic correlation coefficient, CSA- cell surface area, RAW- right atrial weight, RVW-right ventricle weight, LAW-left atrial weight, THW- total heart weight, LVW- left ventricle weight, HR- heart rate, EF-ejection fraction, FS- fractional shortening. All weights are normalized to body weight on day21 and Δ represents difference of measurement from baseline (day21-day0). Purple – common sub-trees in two dendrograms with same hierarchical clustering, green-common sub-trees in two dendrograms with different hierarchical clustering

**Fig. 2.**
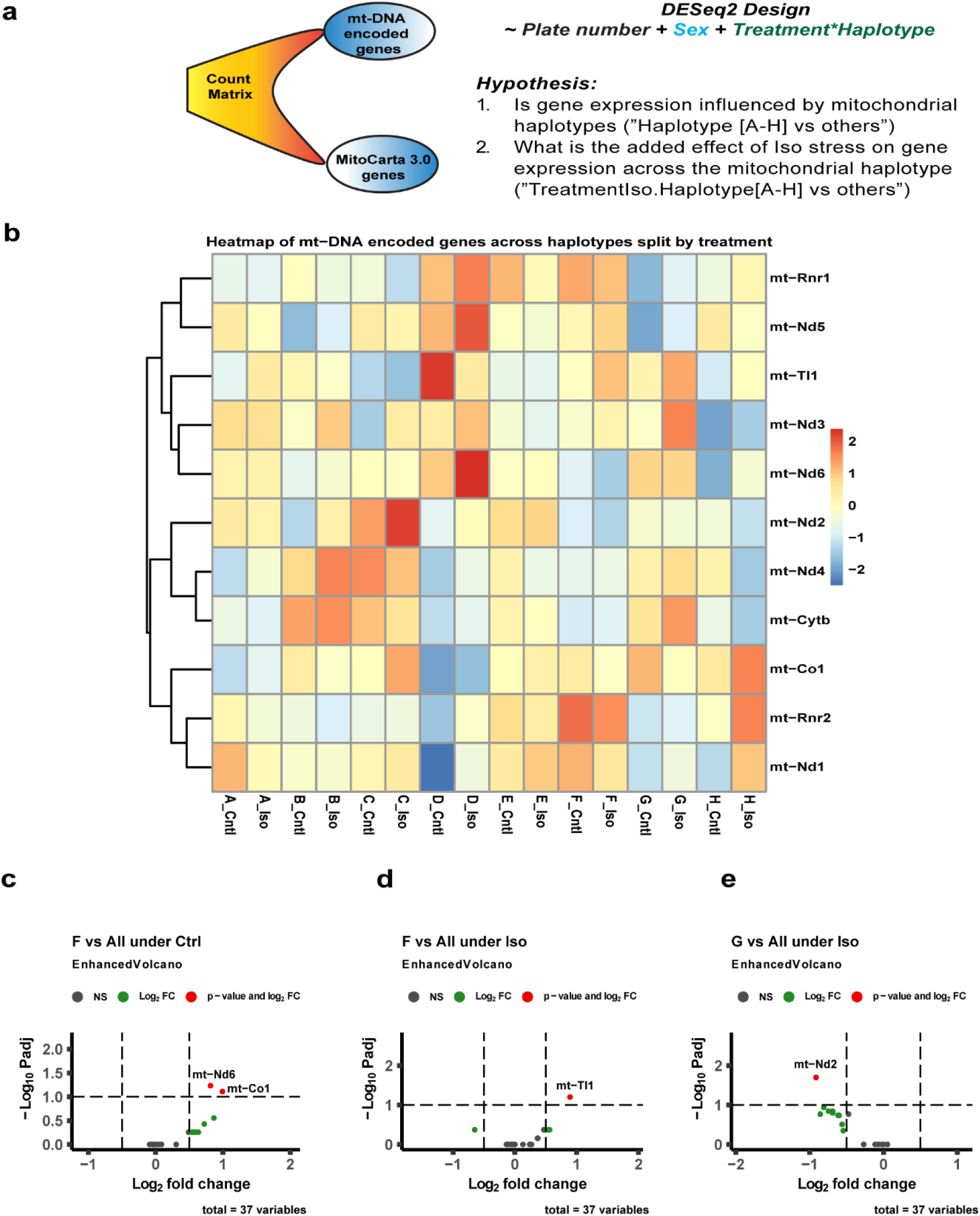
A) Transcriptome data was tested for differentially expressed genes (DEGs) based on the DEseq2 testing design and hypothesis. B) A heat map of expressed (≥ 5 counts in at least 5% of samples) mt-DNA encoded genes is presented across the haplotypes and split by treatment. The significant (padj < 0.1 and log2FC > 0.5) DEGs observed for mt-DNA encoded genes in the haplotypes are presented for c) control groups and d), e) Iso groups. Cntl- control, Iso-isoproterenol, NS- not significant.

For static traits (Fig. 1a), significant (p < 0.05) haplotype differences (haplotype-X vs all other haplotypes in Ctrl/Iso) were noted for mice under control (@:A and B) and Iso (#:A, D, F, G, and H) groups. We observed a higher basal %fibrosis and left ventricular weight (LVW) in mice from haplotypes-A and B respectively in the control arm. On the other hand, the indicated haplotypes from Iso stressed mice had higher %fibrosis (A and H), right atrial weight (RAW: F and G), and total heart weight (THW: G). Interestingly, the weights of total heart (THW), left and right ventricle (LVW and RVW) for haplotype-D and the cell surface area (CSA) for haplotype-F were significantly lower post stress compared to the rest of haplotypes. Treatment differences (Iso vs Ctrl) among each haplotype showed significant (p < 0.05) increase in the Iso-treated cohort for CSA (A), THW (H) and LVW (H).

The dynamic traits (Fig. 1b) showed significant haplotype differences within the control (B) and Iso (C,E,F,G) groups. Left ventricle mass (ΔLV.Mass.Corrected) was the only trait found significantly higher in control mice from haplotype-B. A significant elevation in dynamic traits following Iso stress was noted for haplotype-E (Δ LV.Mass.Corrected, Δ heart rate, Δ ejection fraction and Δ fractional shortening) and haplotype-F (Δ LV.Mass.Corrected) in our study. Treatment differences (Iso vs Ctrl) in dynamic traits were insignificant among these haplotypes.

We next tested to see if similarities across traits matched up with similarities seen across mt-DNA sequences by performing hierarchical clustering of both phenotypic traits and mt-DNA polymorphisms as calculated by ClustalW (see Supplementary figure 1). We explored both static and dynamic traits (Fig. 1c and 1d) using the tanglegram R package [CITE], which measures relationship between two dendrogram clusters. Here we sought to explore to what extent the mt-DNA sequences from the 8 haplotypes influenced cardiovascular traits.

The **static traits** (Fig. 1c) of the 8 mitochondrial haplotypes showed no meaningful relationship (highly tangled) with their founder mt-DNA sequences in the Iso group as evident from high entanglement score and low confidence in their correlation strength (ES:0.905, CPCC: 0.054). This suggests that **remodeling of these static traits is not tightly dependent on mitochondrial sequence variation**. By contrast, the **dynamic traits** (Fig. 1d) exhibited a significant drop in entanglement score(ES = 0.398) and a substantial increase in the strength of correlation (CPCC = 0.444) in the Iso group, suggesting that sequence variation in mt-DNA is superiorly linked to cardiac performance than static traits under adrenergic stress. We did not observe such robust relationship with control mice in either case (Static: ES: 0.501, CPCC: −0.054; Dynamic: ES: 0.787, CPCC: - 0.064). Our results suggest that the mt-DNA sequence association with cardiac performance (dynamics) is dependent on exposure to stress in this case Iso.

### Uniform β-adrenergic stress response of mt-DNA encoded genes across haplotypes contrast with variable expression of nuclear MitoCarta genes

We next explored whether mt-DNA variation between haplotypes drives expression differences between haplotypes and treatments. DESeq2 [CITE] was used to derive differentially expressed genes (DEGs) on two subsets of genes: mt-DNA encoded (37) and from the MitoCarta3.0 database of nuclear genes with strong mitochondrial localization (1140) separately using the design presented in Fig.2a.

Overall gene expression patterns suggested that Iso-induced differential gene expression followed a uniform pattern across haplotypes for mt-DNA encoded genes as we noticed no significant differences when testing for effect of Iso in our design (Difference in expression of Iso vs Ctrl for Haplotype [A-H] vs all other haplotypes). The pattern for mt-DNA encoded genes expressed at a reasonable level (≥ 5 counts) in at least 5% of samples is presented in Fig.2b. However, we did find a total of 20 DEGs from the MitoCarta3.0 list of which twelve were in haplotype-D, four in haplotype-C, two in haplotype-F and one each from E and G haplotypes (Fig.3f). The genes with at least two-fold change in expression included both upregulated genes; *Chdh, Acsm3, Slc25a48, Nat8l, Cyp11b1 and Ucp1* and downregulated genes; *Sdhaf3, Xpnpep3, Abhd10, Slc25a37,Cyp11b1 and Nthl1*.

Next, DEGs for individual treatment groups across the haplotypes were tested (Fig.2c-2e and 3a-e). The F-haplotype had the most DEGs from both mt-DNA and MitoCarta3.0 lists. For mt-DNA encoded genes, *mt-Nd6*, and *mt-Co1* in Ctrl group and *mt-Tl1* in the Iso group were upregulated (padj<0.1 and log2FC > 0.5) in the F haplotype compared to all others (Fig.2c and 2d). Among the nuclear-encoded genes, *Cyp27b1,Fabp1,and Cps1* were upregulated (padj<0.05 and log2FC > 1.0) in the Iso group for the F haplotype compared to all others (Fig.3e). *mt-Nd2* was the only gene found to be downregulated in the G-haplotype compared to the rest of the Iso group (Fig.2e). The remaining haplotypes having differential expressions for nuclear-encoded genes are presented in fig.3a and fig.3b for Ctrl and fig.3c-e for Iso. A heatmap of the gene expression for the 19 MitoCarta genes showing significant difference (padj<0.05 and log2FC > 0.5) under ***stress*** is presented in Fig.3f. These genes, which included (*Ucp1, Sco1, Shmt2, Cyp11b1 and Slc25a37*) were linked to cardiac defects and were enriched for mitochondrial pathways to maintain homeostasis by affecting metabolism of lipids, amino acids, and proteins besides mitochondrial transport, dynamics, and complex activities (Fig. 3g).

**Fig. 3.**
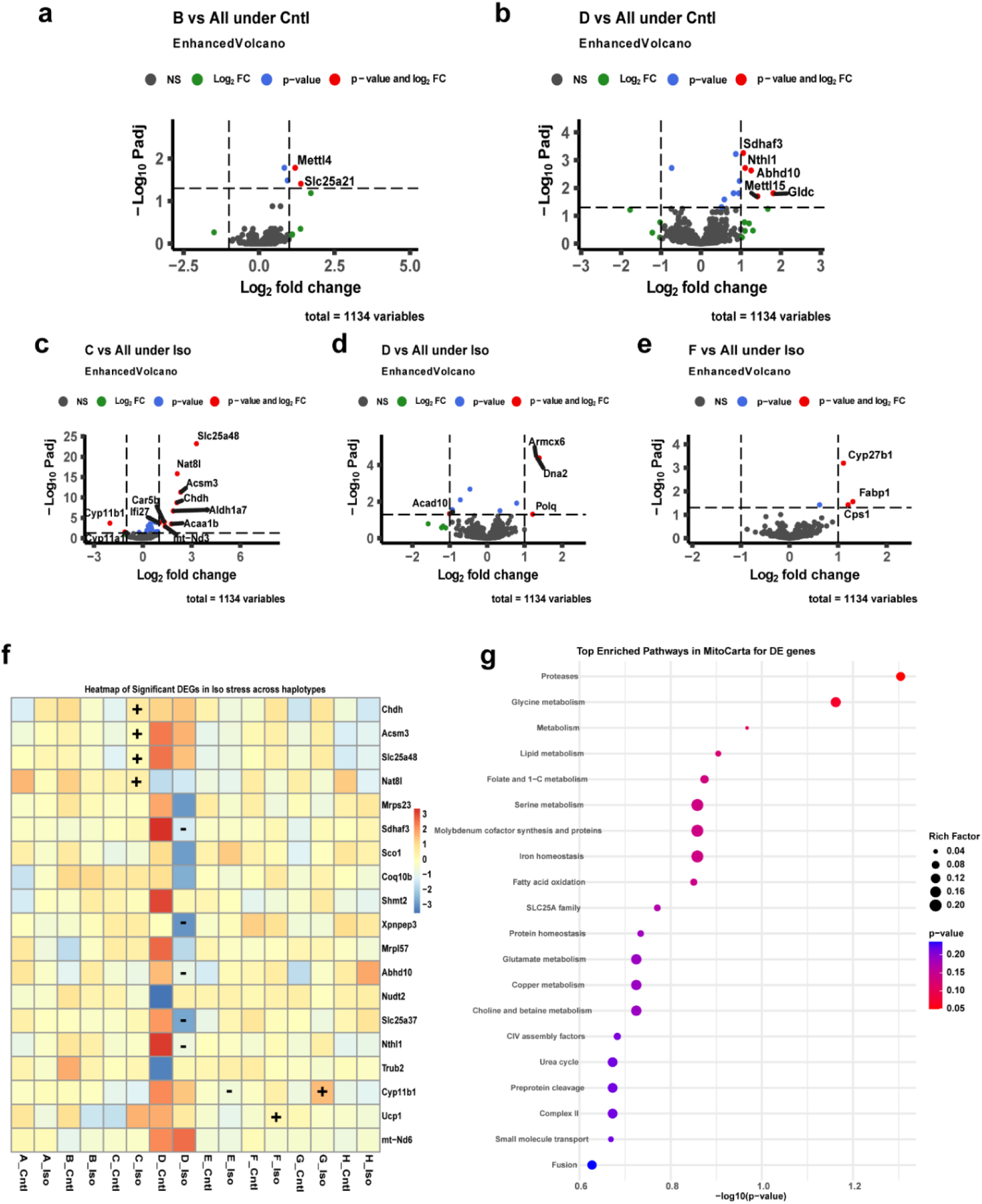
Mitocarta genes were tested for differential expression across haplotypes and significant DEGs are presented in volcano plots for control groups (a, b) and Iso groups (c,d, and e). The added effect of Iso stress on the expression of MitoCarta genes is presented as f) heat map representing the two-fold upregulated genes by ‘+’ and downregulated genes by ‘-’ in their respective haplotype cells across all the haplotypes. Cntl- control, Iso- isoproterenol, NS- not significant.

### GWAS reveals novel loci associated with mitochondria and its nuclear gene expression for heart failure traits

The insights from above suggest that mitochondrial genes (both mt-DNA and nuclear encoded) play a role in shaping cardiovascular dynamic remodeling and are influenced by adrenergic stress. However, a closer link of specific mutations to specific traits was desired. Typically, GWAS approaches are adopted to identify associations of traits with variants in nuclear genes/loci. Largely, such associations fail to identify the complex association of nuclear-mitochondrial signaling (a form of epistasis) due to exclusion of mitochondrial transcripts and their variants from GWAS analyses. Here we use CC variants from the mitochondria across the founder haplotypes (as recorded on the GigaMuga genotyping chip) to uncover signaling mechanisms and associate traits that are central to cardiovascular pathology.

#### Trait-GWAS

Mitochondrial markers were tested for association with physiological traits from CC strains using a linear model. Significant associations with traits from the Iso cohort (cell surface area, delta ejection fraction, right atrial and ventricular weights, and delta fractional shortening) were detected with mt-DNA polymorphisms (Fig. 4a). These traits were influenced by loci encoding for *mt-Nd1, mt-Nd2, mt-Rnr1* and *mt-Rnr2* (Supplementary table 3). Continuing a trend first detected when looking broadly at haplotype-phenotype associations, mice from control cohort did not display any genome-wide significant trait-SNP associations indicating the absence of any associations with mt-DNA under unstressed conditions at this level of power (Fig. 4b and Supplementary table 4).

**Fig. 4.**
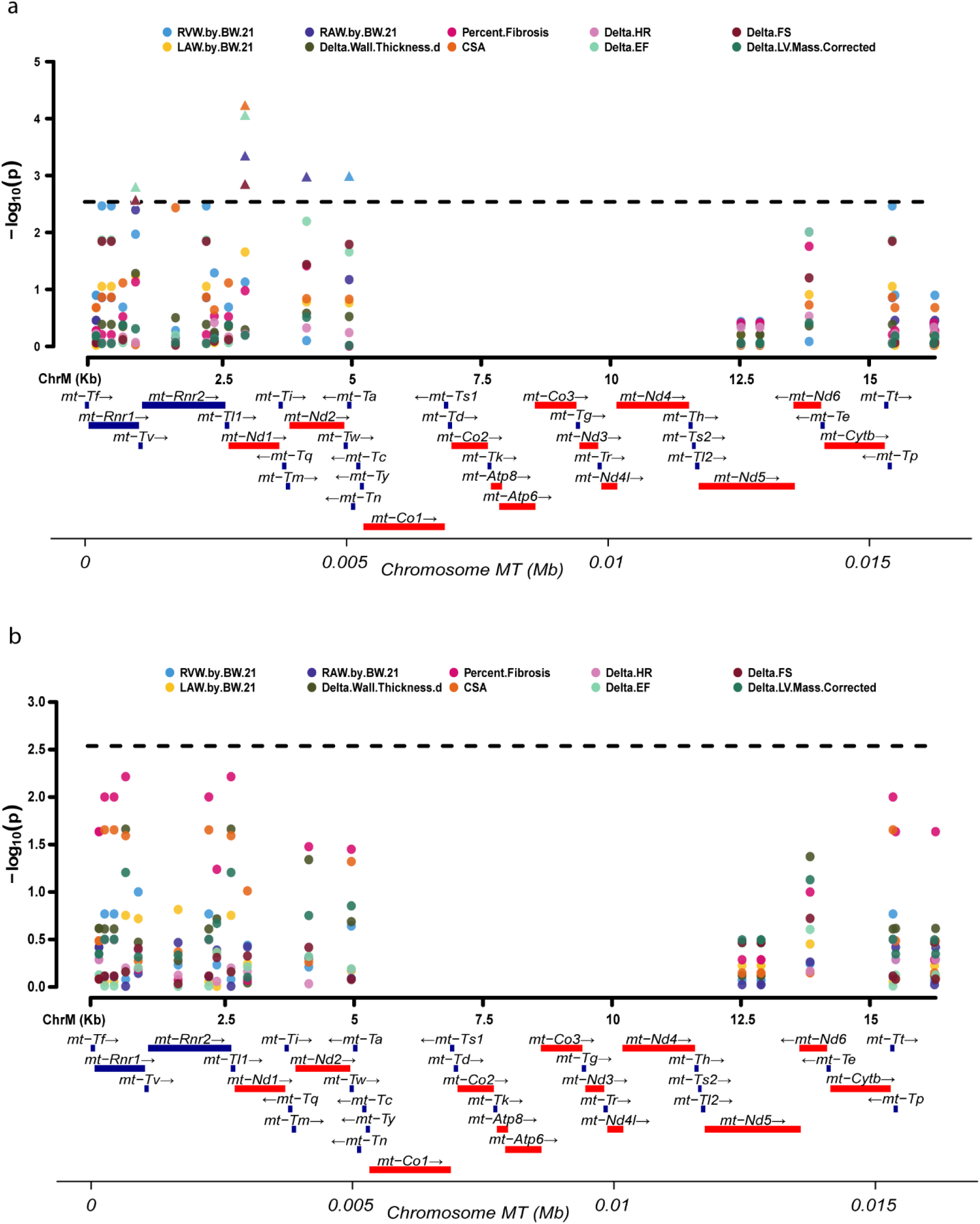
Trait GWAS. a) Isoproterenol group b) Control group. Black dotted line 5% FWER threshold. Gene track red- 13 ETC protein coding genes, blue-rRNA and tRNA encoding regions

#### Trans-eQTL

The trait-GWAS results indicated that there is a polygenic architecture associated with the mitochondrial gene variants that could be influenced by epistatic interactions. These interactions can be detected and mapped through trans-eQTL analyses, especially in the context of mitochondrial-nuclear crosstalk.

Nuclear transcripts from the CC-strains were tested for association with the mitochondrial markers (21 markers from Giga MUGA array). The complete list of gene associations with the mitochondrial markers are provided in supplementary tables 5 and 6. Of 12,439 total genes tested, we observed 831 genes (6.68%) which were significantly associated with one or more marker at a FWER of 5% in the Iso group (Fig. 5d). Of these genes, 47 were present in the MitoCarta gene set and 784 were not thought to localize to the mitochondria (over-representation pValue = 0.57, Fig. 5e). In the control group, we observed 384 genes (3.09%) which are significantly associated with one or more marker at a FWER of 5%. Of these genes, 41 were present in the MitoCarta gene set and 343 were not. This represents a nearly significant (pValue=0.052) over-representation of MitoCarta genes in the control strains (Fig.5e). Fig. 5a shows the top-10 genes associated with markers on the mt-DNA for Iso group. These eQTL associated genes were significantly (p < 0.01) positively enriched for hallmark pathways of oxidative phosphorylation, and negatively for myogenesis, apoptosis, angiogenesis to name a few as represented in fig. 5b with most genes influencing oxidative phosphorylation followed by epithelial to mesenchymal transition (remodeling), coagulation, myogenesis, apoptosis etc., as depicted in fig. 5c.

**Fig. 5.**
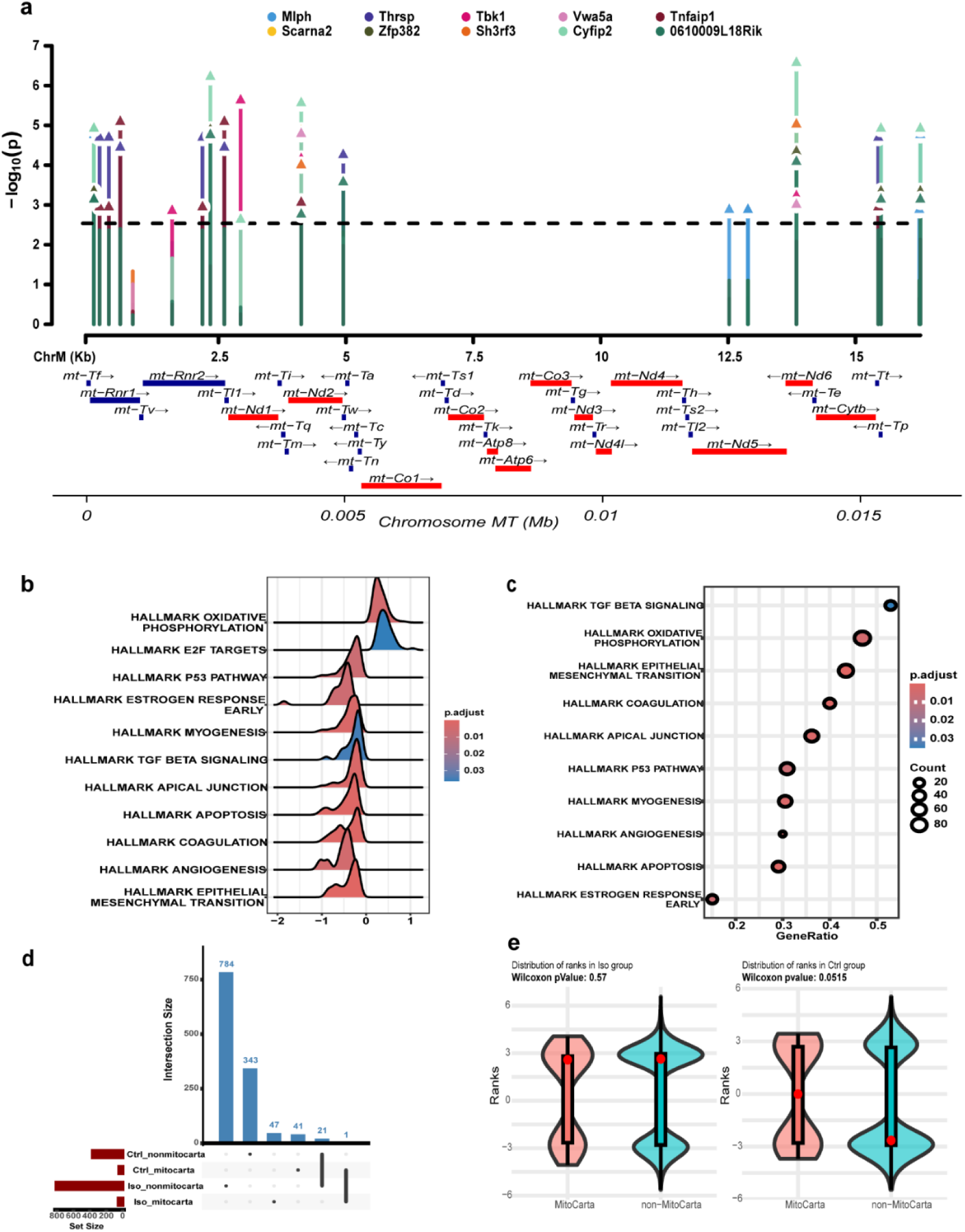
a) eQTL Manhattan plot of top 10 genes in Isoproterenol cohort associated with mitochondrial markers b) Ridge plot of Hallmark pathways enriched in the isoproterenol gene set showing positively and negatively enriched pathways at adjusted pValue < 0.05, c) Dotplot showing pathway represented by maximum genes in the list ordered by gene set size d) Upset plot of significantly associated genes across treatment groups present within MitoCarta and non-MitoCarta subsets e) Overrepresentation analysis for MitoCarta genes based on the ranks from both the treatment groups presented by violin plot and Wilcox-pValues. For d and e, the significant gene list based on 5% FWER threshold was used for plotting.

### Hallmark pathways exclusively observed in Iso-stressed hearts highlight genetic contributors of heart failure

We adopted two approaches to look at genes influencing heart failure. First, the enriched hallmark pathways under Iso stress were assessed for potential complex gene-interactions as depicted by the hallmark pathway networks (Fig. 6a) and critical leading-edge genes were assessed for differences in expression between the treatment groups (Fig.6b) and across the haplotypes (Fig.6c). Second, we overlapped the 831 mt-DNA *trans*-eQTL genes associated with Iso stress with the 648 nuclear-DNA mi-eQTL genes recently reported in our genetic screen of the nuclear genome (CITE). We report these 24 overlapping genes in table 1 as a list of *high confidence gene candidates* associated with potential cardiovascular traits.

**Fig. 6.**
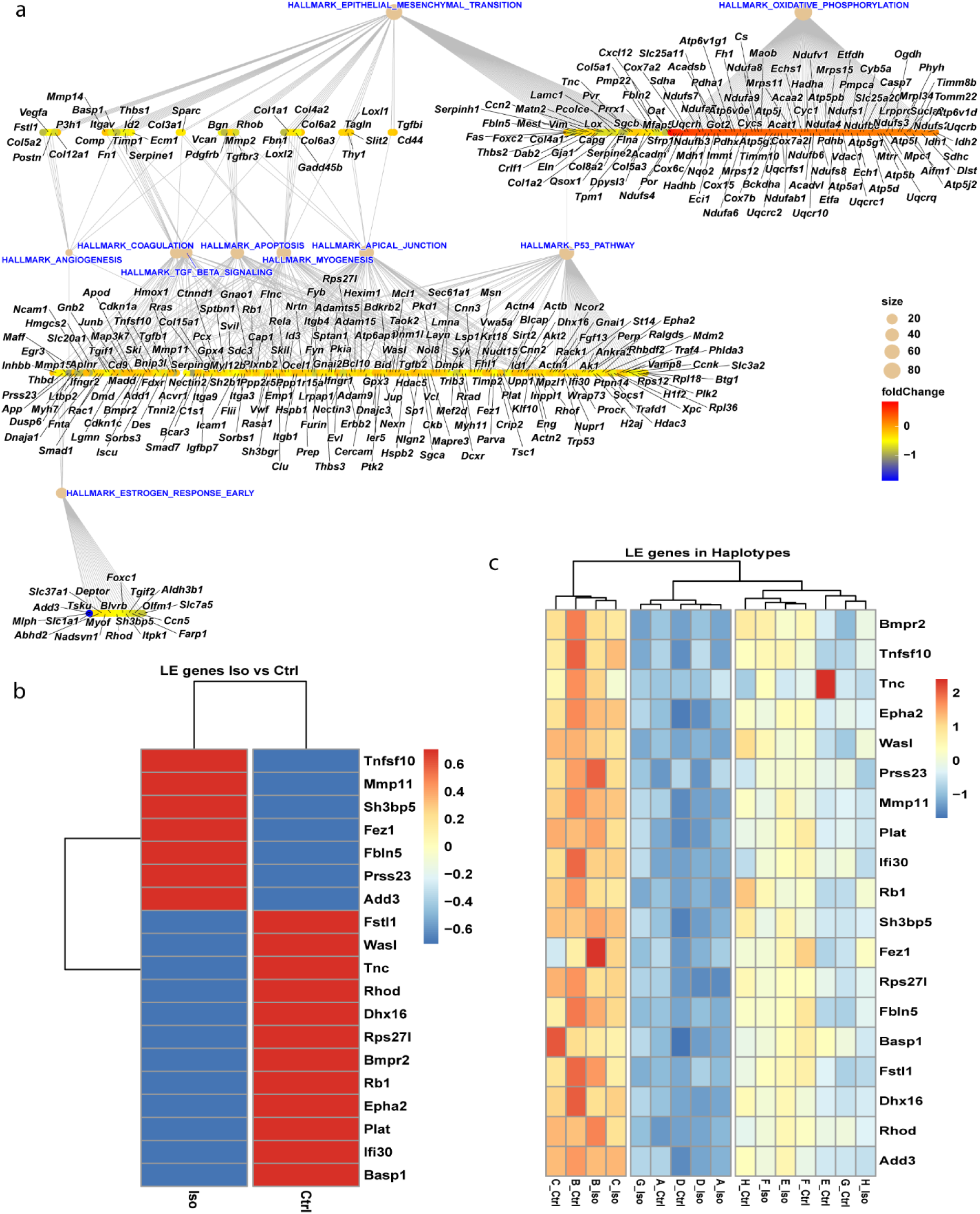
a) GSEA analysis and exploration of gene networks of hallmark pathways showing core enriched genes laid out with Sugiyama algorithm (28) and biological processes associated with significantly enriched pathways in the isoproterenol cohort. Heatmap represents leading edge genes that are significantly (p<0.05) expressed differentially across b) treatments and c) haplotypes.

**Table 1.**
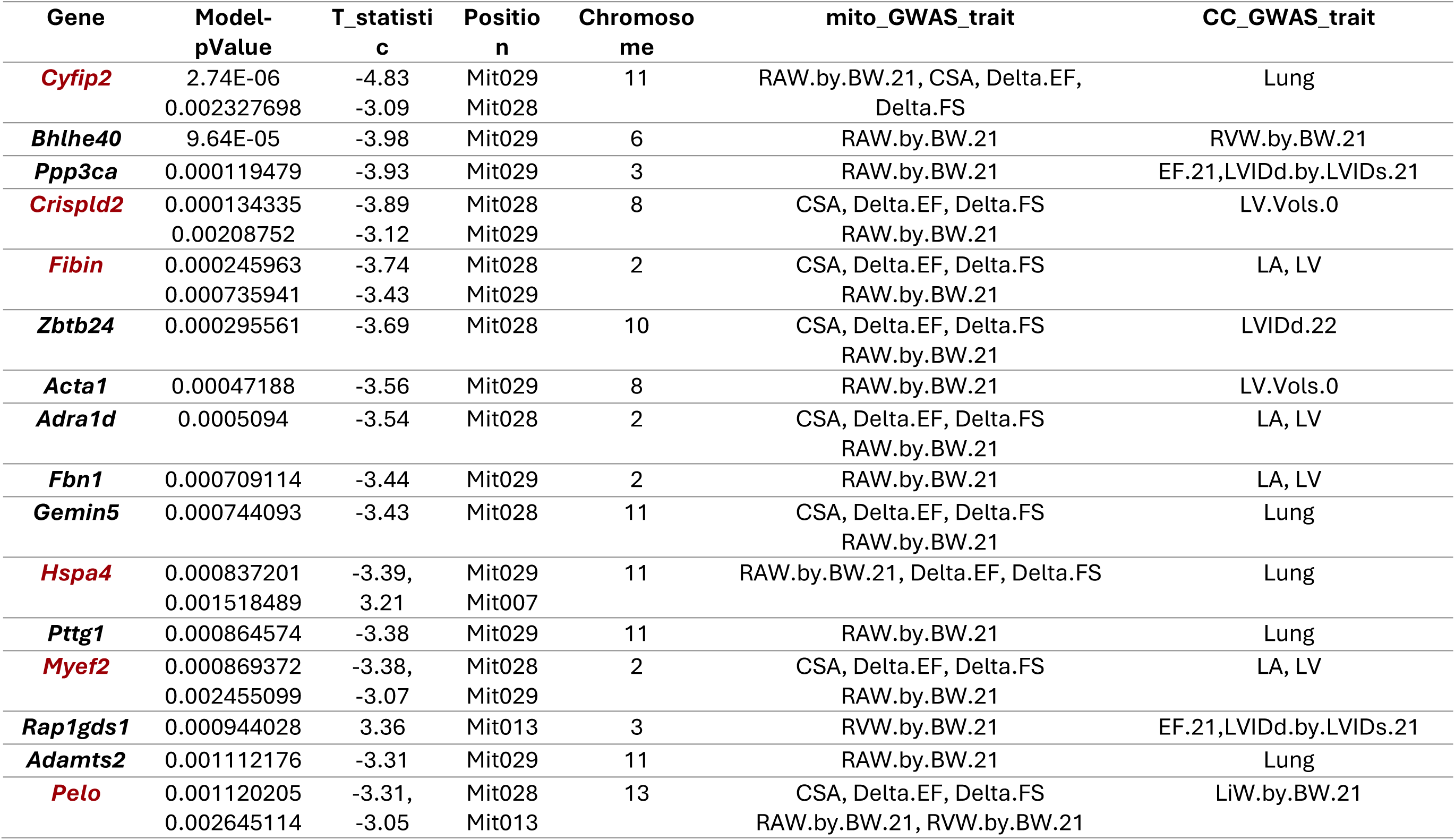
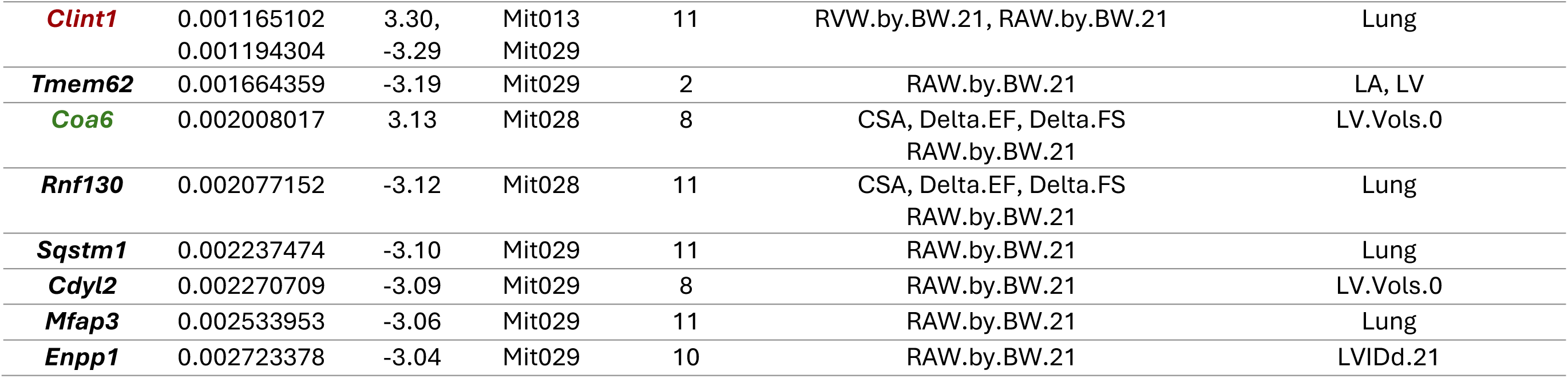
The list of 24 high confidence gene candidates sorted by pValue. Position- position of the eQTL on the mt-DNA, Chromosome-chromosome on which the gene is located, RAW- right atrial weight, RVW-right ventricle weight, CSA-cell surface area, EF- ejection faction, FS- fractional shortening, LA- left atrial weight, LV- left ventricle weight, LVIDd-left ventricle internal diameter at diastole, LVIDs-left ventricle internal diameter at systole, Lung- lung weight. Genes marked in red are associated with multiple positions, gene marked in green is in the MitoCarta3.0 list.(18)

The genes from our eQTL analysis had high positive enrichment only for Oxidative phosphorylation (NES = 1.57,padj = 0.0033) and high negative enrichment for epithelial to mesenchymal transition (NES = −2.56, padj = 2.91e-13) followed by other pathways of interest (supplementary table 7). The leading-edge genes from all the significant pathways were tested using a linear model for expression differences (p < 0.05) across treatments (Fig.6a) and haplotypes (Fig.6b) accounting for differences in both sex and batch effects that arise due to library preparation carried out in five different 96-well plates. Strikingly, we find that mt-DNA haplotype is a stronger predictor gene expression for these genes than the effects of Isoproterenol. We found no hallmark pathways associated with the trans-eQTL genes of control group (results not provided).

Table 1 presents the genes with significant nuclear mi-eQTLs that overlapped with significant mt-DNA trans-eQTL loci. Among the 24 gene candidates, *Cyfip2 returned as top candidate (p =* 2.74e-06 and T = −4.83). These genes are mainly associated with positions Mit007, Mit013, Mit028 and Mit029 of the mt-DNA. *Coa6* (p = 0.002, T = 3.131) was the only gene in the list that was present in MitoCarta, the pathogenic variants of which result in unstable complex-IV subunit seen in patients with neonatal hypertrophic cardiomyopathy (27).

## Discussion

We recently reported a genome scan for novel regulators of heart failure in the Collaborative Cross (CC), a murine genetic reference population (18). In that manuscript, we validated the role of three genes, *Abcb10, Mrps5*, and *Lmod3* on cardiac phenotypes. Each of these genes interact with mitochondria (29–31). In this manuscript, we leverage the ancestry of the CC to better understand the coordinated actions of the mitochondrial (mt-DNA) and nuclear (n-DNA) genomes on the heart. The CC was derived from 8 distinct mouse strains that were bred together such that each strain contains a roughly equal, and randomized, assortment of nuclear genetic material from each of the 8 founders. The *mitochondrial* genetic material, on the other hand, passed down maternally and without recombination, retains the genetic mutations that existed in the original strains, including those between the *domesticus, musculus,* and *castaneus* subspecies from which the parental strains originated. This allows us to ascertain whether changes in mt-DNA, either independent of n-DNA or in conjunction with specific n-DNA variants, lead to changes in heart phenotypes in either healthy or pathological states.

We found that stress-induced functional changes in the heart of CC mice are moderately influenced by founder mt-DNA haplotype, which agrees with prior research showing that CVD risk is associated with mitochondrial haplogroups and variants in humans and mice (5, 32–36). Further, heart failure (HF) being a progressive disease, is controlled by distinct molecular mechanisms, evidenced by structural changes and biomechanical alterations (37). This inspired us to evaluate the HF traits by distinguishing them into two categories (static and dynamic traits). In addition, mitochondria not only provide the energy required for mechanical function but also aids in sustaining the metabolic pathways, both of which have discrete consequences. Haplotypes A and H showed higher % fibrosis and increased cell cross sectional area (CSA) (Fig. 1a and 1b) after Iso, with no signs of cardiac functional decompensation. Haplotype-B had higher heart weights than other haplotypes, while haplotype-D had lower heart weights. Dendrogram analysis (Fig. 1c and 1d) reveals a complex set of relationships between mt-DNA and phenotypes and bolsters these global observations. For example, we observe that strains which contain mitochondria from founder strains F & G entirely converge for static traits irrespective of treatment differences (Fig. 1c). As F & G mitochondria are quite similar to one another, this suggests that this mt-DNA relatedness is resulting in similar phenotypic patterns (the n-DNA from the F & G lines being scrambled across the F & G mitochondrial lines and cancelling out their contributions). We observe that mitochondrial gene expression also aligns with this pattern (Supplementary fig. 2a, ES = 0.288). In contrast, the A and D lines, which also had similar mt-DNA haplotypes in the founder lines, did show some divergence of phenotypes, suggesting possible epistatic interactions and are supported by divergence in the gene expression similarity dendrogram for these haplotypes (Supplementary fig. 2a). Curiously, the dynamic traits (Fig.1d) did not show these same convergences and divergencies, suggesting a greater phenotypic plasticity (remodeling) influenced by physiological state that is conceivably controlled by nuclear-mitochondrial interaction and environmental factors. Evidence for this could be noticed from convergence of mitochondrial gene expression in CC-strains from haplotypes A, D & G for Iso, and Ctrl groups (Supplementary fig. 2d). Unlike static traits, dynamic traits require rapid ATP turnover and calcium handling that depends on the cross-talk between the cytoplasmic and mitochondrial compartments. Hill et.al, demonstrated this reciprocal relationship through stress testing of cardiomyocytes that resulted in loss of respiratory capacity, leading to cell death (38). In support, we noticed that haplotypes A, D & G display a more pronounced trait spectrum under stress (for example we observed a 12-14% increased CSA to be accompanied by 27-33% higher fibrosis in A and G, while a 14% increased CSA resulted in <1% increase in fibrosis in D) compared to the rest of the haplotypes.

We next decided to look deeper at the individual polymorphisms that exist within the mt-DNA and their relationship with HF-associated phenotypes. Mitochondrial-GWAS (Mito-GWAS) is traditionally challenging, with weak or non-existent signals due to their unique maternal inheritance pattern, heteroplasmy, small number of variants in the 16Kb genome and interaction with – and overshadowing by - the nuclear genome. The CC offers a unique opportunity to perform Mito-GWAS due to the way it was constructed, which scrambles and masks the contributions of the nuclear genome and allows us to use a linear mixed model to link phenotypes to mt-DNA polymorphisms. We observe significant associations (pValues less than the 1% Family-Wise Error Rate (FWER) of 0.0029) between four sites (*Mit007, Mit013, Mit028 and Mit029* at positions 817, 2934, 4123 and 4947, respectively) and five phenotypic traits: change in ejection fraction, change in fractional shortening, cardiomyocyte cross sectional area, right ventricular weight and right atrial weight) in Iso treated animals (Fig. 5a). Genes at these sites are responsible for energy production (*mt-Nd1, mt-Nd2)* and ribosomal function (*mt-Rnr1)*. Mutations in these genes have been implicated in mitochondrial diseases such as LHON, MELAS and metabolic disease that can lead to accelerated cardiovascular disease progression. In 2020, a non-pathogenic variant in m.G14904A (mt-Cyb) was identified as one such modifier for disease phenotype in a *Bcs1l ^p.S78G^* that reduced the lifespan and exacerbated the disease progression in mice demonstrating mito-nuclear epistasis (39).

Since the primitive bacteria that would eventually become the mitochondria first entered into a symbiotic relationship with the larger cell that would be its host, the mitochondria have progressively outsourced many of their genes that make up its protein complement to the genome as nuclear-encoded mitochondrial transcripts. These nuclear genes and the genes retained on the mitochondrial genome are in constant communication with one another. We were interested in exploring whether specific mt-DNA haplotypes resulted in changes in these relationships in both healthy and stressed hearts. To do so, we utilized the MitoCarta3.0 database (23) to extract all mitochondria-associated genes from our CC transcriptomes, and also examined changes in gene expression for the explicitly mt-DNA-coded genes as well. Several compensatory mechanisms stabilize heart function that act through mitochondria to ensure that high energy demands are met even under stress (19, 40). However, under prolonged stress, (in our case 21 days of continuous adrenergic stimulation) decompensating conditions worsen the crisis and lead to heart failure. Studies have shown that in such conditions, increased expression of *Ucp1* protects the heart through preserving mitochondria and reducing oxidative stress (41, 42). We observed several such genes activated in response to stress (Fig. 3f). Clearly, in the case of haplotype-F, upregulation of *Ucp1* (log2FC = 2.03, baseMean = 10.15) was noted along with elevated expression of mt-DNA encoded genes (*mt-Tl1, mt-Nd6 and mt-Co1*) in these mice. As expected, these mice showed lower cell surface area (hypertrophy), and fibrosis following Iso stress. On the contrary, upon examining the haplotype-B which had lower *Ucp1* expression under stress, we noted a higher damage to the myocardium (Fig.1a). Although the tests indicate no significant change from control, we suspect this could be due to variation within the strains of the haplotype-B as the gene expressions poorly correlate (Supplementary fig 2d). Consistent with our findings, some of the DEG’s found in this study (*Sco1, Shmt2, Cyp11b1 and Slc25a37*) are reported as an underlying cause for cardiac defects (43–47). Thus, cardiac manifestations in these haplotypes could be linked to nuclear genetic defects due to mitochondrial disorders. While a direct link of some of these gene is still under investigation, the differential expression of these genes in certain haplotypes of CC-strains could benefit choosing them as models for such studies. It is important to note that though the expression changes across the haplotypes seem to not affect the heart under normal physiology, there is a high risk/susceptibility of developing heart failure under stress. For instance, although trait differences existed across the haplotypes within each treatment, only haplotypes **A** and **H** (indicated by * in Fig.1a) showed significant difference (Iso vs Ctrl) in total heart weight, LV weight and CSA when subjected to stress. Ballinger’s group has demonstrated that Mito-nuclear genetic interactions are known to affect gene expression in obesity but their precise mechanism are under investigation (48).

Next, we integrated our Mito-GWAS and transcriptome approaches together to perform *trans*-eQTL analyses of n-DNA genes with mt-DNA polymorphisms. We observe 831 associated (pValues less than the 1% FWER of 0.0029) n-Genes with mt-DNA SNPs in Iso-treated samples. The top 10 candidates (p < 2.02e-05) had association with mitochondria and cardiovascular disease (Fig. 6a). *Tbk1* deletion from cardiomyocytes aggravates cardiotoxicity by inhibiting mitophagy (49). *Sh3rf3* is a potential regulator of hypertrophy and fibrosis by controlling fibroblast-myofibroblast differentiation (50)*. Cyfip2* is implicated in neuronal function by altering the mitochondrial content and the KO mice showed congenital heart defects (51, 52). Inhibition of *Tnfaip1* protected cardiomyocytes from ischemia-reperfusion injury (53). *Thrsp* plays a critical role in fatty acid metabolism, and silencing the gene impaired mitochondrial respiration in adipocytes (54). Recently, a study on Chinese population showed *Mlph* hypermethylation in blood of patients with coronary heart disease (*55*). Lower *Scarna2* levels were noted in congenital heart defect cases and altering the levels of these non-coding RNAs were lethal to embryo due to faulty mRNA processing as these genes are involved in mRNA splicing (56). The remaining genes in the top 10 most associated ones *(Vwa5a,Zfp382 and 0610009L18Rik)* are all expressed in the heart but their functional role in cardiac phenotypes are not known at this time.

Analysis of the larger gene list for effects on major hallmark pathways revealed oxidative phosphorylation as the only hallmark pathway which showed positive enrichment while many others were negatively enriched, including genes involved in the epithelial-mesenchymal transition, angiogenesis, apoptosis, the apical junction, myogenesis, TGF beta signaling, coagulation and early estrogen response (Fig.5b and Fig.6a). *Atp6v0e* and *Cyb5a* were the only genes controlling OXPHOS that were eQTL hits found in MitoCarta3.0, an indication that nuclear genome influences major pathways in a stressed myocardium. *Atp6v0e* is a component of ATP dependent proton pumps (V-ATPase) in the heart that regulate intracellular pH, disruption of which can lead to heart failure. In a study where ATP6AP2, another component of the pump, was knocked out, heart function deteriorated due to dysfunctional autophagy leading to mitochondrial dysfunction (57). There is no direct evidence linking the *Cyb5a* gene to heart failure although it does act as an electron carrier for *Cyb5r3* a known contributor to heart failure and so presumably it is linked to heart failure through a similar pathway (58).

Twenty-four candidate genes present within our n-DNA GWAS study loci overlap with mt-GWAS trans-eQTL hits as summarized in table 1. Among these genes are five well-known genes associated with cardiac hypertrophy (*Fibin*, *Hspa4*)(59, 60) and heart failure (*Enpp1*, *Adamts2*, *Sqstm1*)(9, 61–63). Additionally, all these genes are involved in maintenance of mitochondrial function directly or as mitokines during stress as reported by multiple studies (64–69). *Coa6* is the only candidate directly involved with complex-IV assembly, mutations of which have been reported in cardiomyopathy cases (70). It is critically involved in biogenesis of mt-DNA encoded COX2 but its role in HF remains to be explored.

Returning to the haplotypes, we examined how gene expression patterns for our 24 candidate eQTL genes varied across the haplotype groups during heart failure progression (supplementary figure 3). For instance, *Pttg1, Cyfip2, and Acta1* were noticeably downregulated in haplotype-A, the mice from which had significantly higher %fibrosis & increased CSA following Iso challenge, matching their proposed roles of fibroblast activation and proliferation (71–73). Likewise, mice in haplotype-E showed a higher expression of these genes and a lower increase in cardiac fibrosis and higher functional performance post Iso stress. Interestingly another gene *Cdyl2,* a critical epigenetic regulator of gene expression was visibly upregulated in haplotype-E (74). Yunfei et.al, reported that a circular CDYL2 promotes apoptosis in cardiomyocytes following myocardial infarction in rat model(75). Such contrary reports suggest that potential mitochondrial axis exists that decides the nuclear gene’s expression and needs to be studied in detail. These links between haplotypes and phenotypes, mediated by gene expression, are strong evidence for the role of these different haplogroups of mitochondria, independent of n-DNA, as drivers and mediators of cardiac remodeling during beta adrenergic overdrive scenarios. A greater understanding of the role of these mitochondrial haplotypes through an examination of different phenotypes and diseases in a cohort like the CC could lead to further revelations regarding the role of mitochondrial differences on other known strain-to-strain variations.

In conclusion, in this study we exposed CC strains to experimental heart failure and studied cardiovascular traits and genetics through the lens of the eight mitochondrial haplotypes present within the cohort. Our study highlighted that stress driven cardiac phenotypes may be driven by distinct inherited polymorphisms in the mt-DNA that act either independently or, more often, in conjunction with polymorphisms on n-DNA to alter phenotypic presentations. These results are consistent with prior evidence that suggests that changes in nuclear gene expression can occur in response to modest changes in mitochondrial DNA mutations that can influence progression and severity of heart disease (76, 77). Our present mito-GWAS approach forms the basis for uncovering mito-nuclear interactions that underly any disease condition, and candidate genes identified by this approach hold promise for studying such interactions in the context of HF. Future work using specific CC strains could seek to better disentangle these mt-DNA/n-DNA relationships through genetic manipulation of specific n-DNA/mt-DNA sites via CRISPR (32).

Our work has some limitations, the largest of which is the limited diversity of mitochondrial SNPs and the lack of heteroplasmy information across the strains, with our haplotype markers not fully accounting for every mutation present in the CC mitochondria. This could be addressed by full-length sequencing of the mt-DNA to find strain specific somatic variants and calculating the percentage heteroplasmy, improving the study rigor. We are also limited by the number of strains in the CC, with more strains providing a better ‘mask’ for the role of n-DNA in heart phenotypes when examining the mitochondrial haplotypes. Finally, we would like to see researchers in the field utilize this approach to uncover anterograde-retrograde signaling mechanisms governing mitochondrial-pathologies to address complex trait diseases.

## Acknowledgement

The authors would like to thank the Small Animal Imaging Core Facility, funded in part by the NCI Cancer Center Support Grant, P30CA016086, and the NIH S10 Shared Instrumentation Grant, S10OD034328, for supporting this work.

## Author contributions

Conceptualization and funding: Christoph D. Rau

Methodology: Todd H. Kimball, Sriram Ravindran, Caitlin Lahue and Christoph D. Rau

Data curation: Sriram Ravindran, Brian Gural

Investigation and Project administration: Todd H. Kimball, Caitlin Lahue, Sriram Ravindran and Brian Gural

Writing – Original draft: Sriram Ravindran and Christoph D. Rau

Writing – Reviewing and editing: All authors

